# A novel microtubule nucleation pathway for meiotic spindle assembly in oocytes

**DOI:** 10.1101/284901

**Authors:** Pierre Romé, Hiroyuki Ohkura

## Abstract

The meiotic spindle in oocytes is assembled in the absence of centrosomes, the major microtubule nucleation sites in mitotic and male meiotic cells. A crucial, yet unresolved question in meiosis is how spindle microtubules are generated without centrosomes and only around chromosomes in the large volume of oocytes. Here we report a novel oocyte-specific microtubule nucleation pathway that is essential for assembling most spindle microtubules complementarily with the Augmin pathway, and sufficient for triggering microtubule assembly in oocytes. This pathway is mediated by the kinesin-6 Subito/MKlp2, which recruits the γ-tubulin complex to the spindle equator to nucleate microtubules in *Drosophila* oocytes. Away from chromosomes, Subito interaction with the γ-tubulin complex is suppressed by its N-terminal region to prevent ectopic microtubule assembly in oocytes. We further demonstrate that the Subito complex from ovaries can nucleate microtubules from pure tubulin dimers *in vitro*. Taken together, microtubule nucleation regulated by Subito drives spatially restricted spindle assembly in oocytes.

## Introduction

Spatial and temporal regulation of microtubule nucleation is vital for the formation and maintenance of a functional spindle. In mitotic or male meiotic animal cells, centrosomes are the major microtubule nucleation sites, which appear to be central to define the position of the spindle formation and the spindle bipolarity. Despite the apparent central roles of centrosomes, the bipolar mitotic spindle can be formed in mitotic cells even when centrosomes are artificially inactivated (Khodjakov et al., 2000; Hinchcliffe et al., 2001). Moreover, centrosomes have been shown to be dispensable for the flies, as Drosophila without centrosomes can survive to the adult stage (Basto et al., 2006).

In most animals, female meiosis in oocytes is different from mitosis or male meiosis, as oocytes naturally lack centrosomes to assemble the meiotic spindle (McKim and Hawley, 1995; Karsenti and Vernos, 2001). The lack of centrosomes in oocytes poses two fundamental questions: how are spindle microtubules nucleated, and how is efficient nucleation spatially restricted to only around the chromosomes? A constantly high level of nucleation is essential for forming and maintaining the spindle, due to the intrinsically dynamic nature of spindle microtubules (Mitchison and Kirschner, 1984). In addition, as oocytes are exceptionally large in volume, it is crucial to spatially limit a high level of nucleation only to the vicinity of chromosomes.

In mitotic cells, centrosome-independent microtubule nucleation takes place randomly along the spindle microtubules (Mahoney et al., 2006). This nucleation has been shown to be mediated by a conserved 8-subunit complex called Augmin (Goshima et al., 2008; Meireles et al., 2009; Uehara et al., 2009; Kamasaki et al., 2013). Augmin sits on a pre-existing microtubule and recruits the γ-tubulin complex by direct interaction with one of the γ-tubulin subunits, NEDD1 (Grip71 in *Drosophila*) (Uehara et al., 2009; Chen et al., 2017). The γ-tubulin complex then nucleates a new microtubule from a pre-existing one, leading to “microtubule amplification” which exponentially increases the number of microtubules (Goshima and Kimura, 2010). In mitotic cells, a loss of either Augmin or Grip71 leads to the same dramatic decrease in the spindle microtubule density (Meireles et al., 2009; Reschen et al., 2012), demonstrating that this microtubule amplification pathway is responsible for the majority of microtubule assembly even in the presence of centrosomes. Furthermore, loss of Augmin leads to severe reduction of spindle microtubules in mitotic cells which lack centrosomes (Goshima et al., 2008; Meireles et al., 2009; Wainman et al., 2009), showing that Augmin-mediated microtubule nucleation accounts for the assembly of virtually all centrosome-independent spindle microtubules in mitosis.

Surprisingly, a robust bipolar spindle can be formed without the Augmin complex in *Drosophila* oocytes that naturally lack centrosomes (Meireles et al., 2009). This demonstrates that oocytes have an alternative pathway which can assemble spindle microtubules without both Augmin and centrosomes. In stark contrast to the loss of Augmin, removal of the γ-tubulin subunit Grip71 in oocytes strongly reduces spindle microtubules (Reschen et al., 2012). This paradoxical result strongly suggests the existence of another nucleation pathway specific to oocytes for assembling a meiotic spindle.

Here we report a novel microtubule nucleation pathway in oocytes, which is mediated by a kinesin-6, Subito/MKlp2. We show that Subito and Augmin recruit Grip71 to the spindle equator and poles, respectively. These two pathways act complementarily to assemble most of the spindle microtubules in oocytes. Furthermore, the N-terminal region of Subito is important to spatially restrict the spindle microtubule nucleation only to the vicinity of meiotic chromosomes. Therefore, this novel nucleation pathway is central for both assembling a meiotic spindle around chromosomes and preventing ectopic microtubule nucleation in the large volume of an oocyte.

## Results

### Grip71/NEDD1 is recruited to the spindle equator in the absence of Augmin in oocytes

In mitotic *Drosophila* cells, few microtubules are assembled in the absence of both centrosomes and Augmin (Goshima et al., 2008; Meireles et al., 2009; Wainman et al, 2009). In contrast, in *Drosophila* oocytes which naturally lacks centrosomes, we have previously shown that spindle microtubules are robustly assembled in the absence of Augmin (Meireles et al., 2009). This difference points to the presence of a yet unknown, oocyte-specific microtubule assembly pathway.

Spindle microtubule assembly in oocytes is greatly reduced in the absence of the γ-tubulin subunit Grip71/NEDD1 (Reschen et al., 2012), suggesting that this new oocyte-specific pathway largely depends on Grip71. Grip71/NEDD1 is a γ-tubulin subunit through which Augmin recruits the γ-tubulin complex to existing microtubules (Vérollet et al., 2006; Luders et al., 2006; Zhu et al., 2008; Uehara et al., 2009; Chen et al., 2017). In mitosis, localisation of γ-tubulin and Grip71 on the spindle microtubules depends entirely on Augmin (Goshima et al., 2008; Zhu et al., 2008, Wainman et al., 2009). We hypothesised that oocytes have an alternative Augmin-independent pathway which recruits the γ-tubulin complex onto the spindle microtubules through Grip71.

To test this hypothesis, an antibody was raised against Grip71(**Fig S1**) and used to immunostain mature wild-type oocytes which naturally arrest in metaphase I. We found that Grip71 is concentrated at the spindle poles and equator (**Fig 1**). Next, to test whether this localisation depends on Augmin, Grip71 was immunostained in oocytes of a null mutant of the essential Augmin subunit Wac (Meireles et al., 2009). Concentration of Grip71 at the spindle poles was greatly reduced in *wac* mutant oocytes in comparison with wild type, but Grip71 localisation to the spindle equator was not affected (**Fig 1**). This Augmin-dependent Grip71 localisation to the spindle poles is consistent with our previous observation of Augmin localisation to the poles and its function to promote microtubule assembly near the poles in oocytes (Meireles et al., 2009; Colombie et al., 2013). In contrast, the localisation of Grip71 to the spindle equator in oocytes is independent of Augmin, which can explain why oocytes robustly assemble spindle microtubules without Augmin, but fail to do so without Grip71 (Meireles et al., 2009; Reschen et al., 2012).

**Figure 1.**
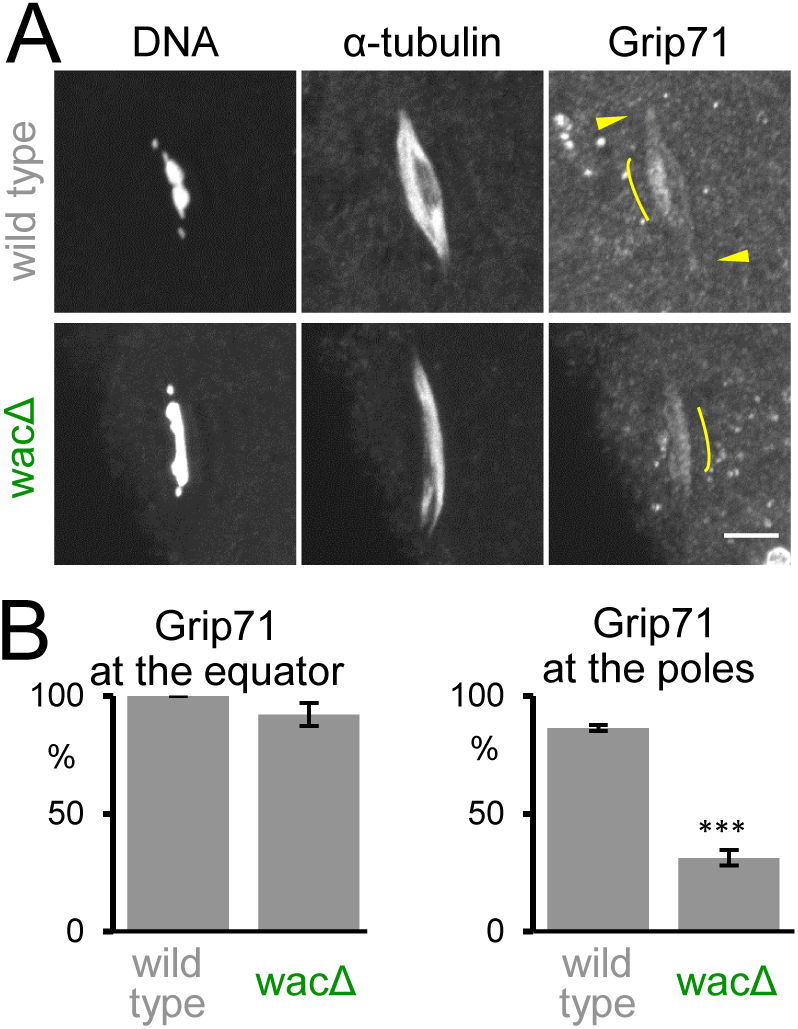
Grip71 is recruited to the spindle equator independently of Augmin in oocytes. (A) Immunostaining of a meiotic spindle in mature oocytes from wild type and a deletion mutant of *wac* gene encoding an essential subunit of the Augmin complex. Arrowheads indicate Grip71 at the spindle pole, and curved lines indicate the concentration of Grip71 in the spindle equator. Pole localisation of Grip71 was greatly reduced in *wac* mutant. Bar=5 µm. (B) The frequencies of meiotic spindles which have Grip71 concentration at the spindle equator and Grip71 localisation at at least one of the spindle poles, respectively. The graphs show the means (the main bars) and standard errors (error bars) from three repeats of experiments for wild type and *wac*Δ (38, 41 spindles in total). *** indicates a significant difference from wild type (p<0.001).

### The *Drosophila* MKlp2, Subito, recruits Grip71 to the spindle equator

To understand the molecular mechanism of this novel microtubule assembly pathway, we sought to identify a protein which recruits Grip71 to the spindle equator. We predict that such a protein should (1) colocalise with Grip71 to the spindle equator in oocytes, (2) physically interact with the Grip71/γ-tubulin complex directly or indirectly, and (3) be required for Grip71 localisation to the spindle equator but not the spindle poles. We considered Subito (the orthologue of mammalian MKlp2, a kinesin-6) to be a good candidate, as it was previously shown to localise to the spindle equator and have an important role in the integrity of the spindle equator (Jang et al., 2005).

To test whether Subito colocalises with Grip71 to the spindle equator, mature oocytes were co-immunostained for Subito, Grip71 and α-tubulin (**Fig 2**). Subito and Grip71 showed nearly identical localisation patterns in the spindle equator, while α-tubulin showed a distinct pattern from them (**Fig 2A**). This was confirmed by intensity measurements along a cross-section of a spindle (**Fig 2A**). To test physical interaction between Subito and Grip71, GFP-Subito expressed in ovaries was immunoprecipitated for analysis by western blot. Grip71 and γ-tubulin were both co-immunoprecipitated with GFP-Subito from a *Drosophila* ovary extract (**Fig 2B**), demonstrating that Subito interacts with Grip71 and γ-tubulin.

**Figure 2.**
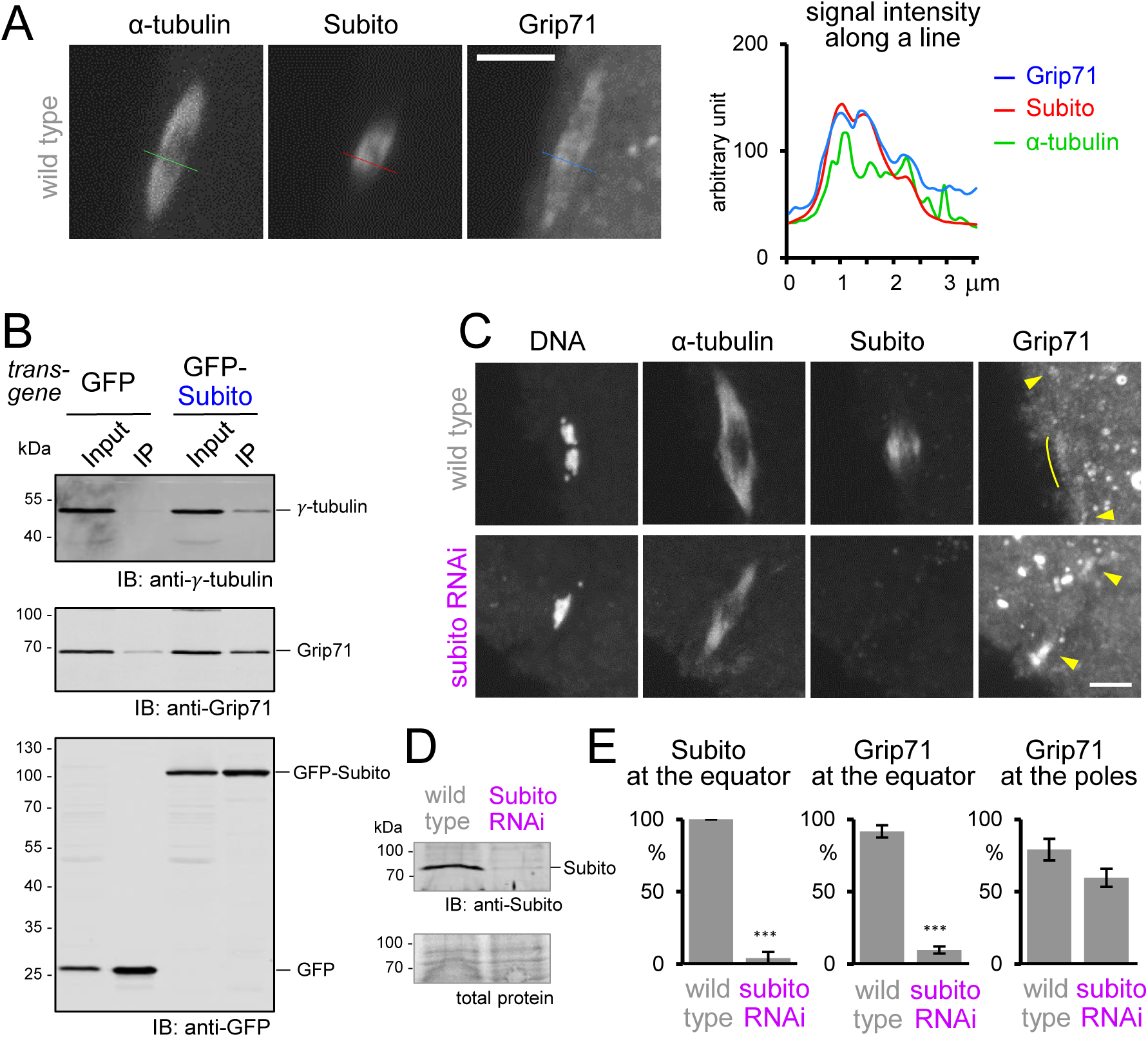
Subito, the *Drosophila* MKlp2, recruits Grip71 to the spindle equator. (A) Colocalisation of Subito with Grip71 in the spindle equator. Immunostaining of a meiotic spindle in mature wild-type oocytes, and signal intensities (arbitrary unit) along the line. Grip71 and Subito are correlated with each other, better than with α-tubulin. Bar=5 µm. (B) Subito physically interacts with γ-tubulin and Grip71. GFP or GFP-Subito was immunoprecipitated from oocytes expressing these proteins in the absence of phosphatase inhibitors, followed by immunoblots (IB). The figure shows one of biological triplicates, all of which showed co-immunoprecipitation. 10% of Input relative to the immunoprecipiated fraction (IP) was loaded. (C) Subito is required for Grip71 localisation to the spindle equator. Immunostaining of mature wild-type and *subito* RNAi oocytes. Arrowheads indicate Grip71 localisation at the spindle pole, and curved lines indicate Grip71 concentration in the spindle equator. The equator localisation of Grip71 was lost in the *subito* RNAi oocyte. Bar=5 µm. (D) Immunoblot of ovaries from wild-type and *subito* RNAi flies. (E) The frequencies of meiotic spindles which have Subito and Grip71 concentration at the spindle equator and Grip71 localisation at at least one of the spindle poles, respectively. The graphs show the means and standard errors from three repeats for each of wild type and *subito* RNAi (86, 61 spindles in total). *** indicate significant differences from wild type (p< 0.001, respectively).

To test whether Grip71 localisation depends on Subito, mature oocytes expressing short hairpin RNA (shRNA) against *subito* were immunostained for Subito, Grip71 and α-tubulin (**Fig 2C**). Subito signals were dramatically reduced, confirming the specificity of the anti-Subito antibody and the effectiveness of RNAi (**Fig 2C-E**). Strikingly, Grip71 localisation to the spindle equator was lost (**Fig 2C,E**). In contrast, Grip71 localisation at the poles was still observed, although in fewer oocytes (**Fig 2C,E**). This shows that Subito is essential for Grip71 localisation to the spindle equator, but largely dispensable for its localisation to the poles. Taken together, we conclude that Subito recruits Grip71 to the spindle equator in oocytes.

### Subito and Augmin complement each other to generate spindle microtubules

It was previously shown that the total amount of spindle microtubules was not significantly reduced in a null mutant of either Subito or the Augmin subunit Wac in oocytes (Giunta et al., 2002; Jang et al., 2005; Colombié et al., 2008; Meireles et al., 2009). We hypothesised that these two pathways may act complementarily to assemble spindle microtubules. To test this, both Subito and Wac were simultaneously depleted from oocytes by expressing short hairpin RNA (shRNA) against Subito in ovaries of the *wac* null mutant. Immunostaining showed that a single depletion of Subito or Wac did not reduce the total amount of spindle microtubules (**Fig 3C**), as previously reported (Giunta et al., 2002; Jang et al., 2005; Colombie et al., 2008; Meireles et al., 2009). In contrast, simultaneous depletion of Subito and Wac led to a dramatic reduction of spindle microtubules and often thinner spindles (**Fig 3A,B**). This phenotype is reminiscent of the *grip71* RNAi or null mutant phenotype (**Fig 3A,B**; Reschen et al., 2012), suggesting that Grip71 recruitment to the meiotic spindle depends entirely on Subito and Wac. Indeed, immunostaining showed that Grip71 fails to localise to the spindle in oocytes co-depleted of Subito and Wac (**Fig 3A,C**). Therefore, we conclude that Subito and Augmin mediate two complementary pathways which recruit Grip71/γ-tubulin to assemble most spindle microtubules in oocytes.

**Figure 3.**
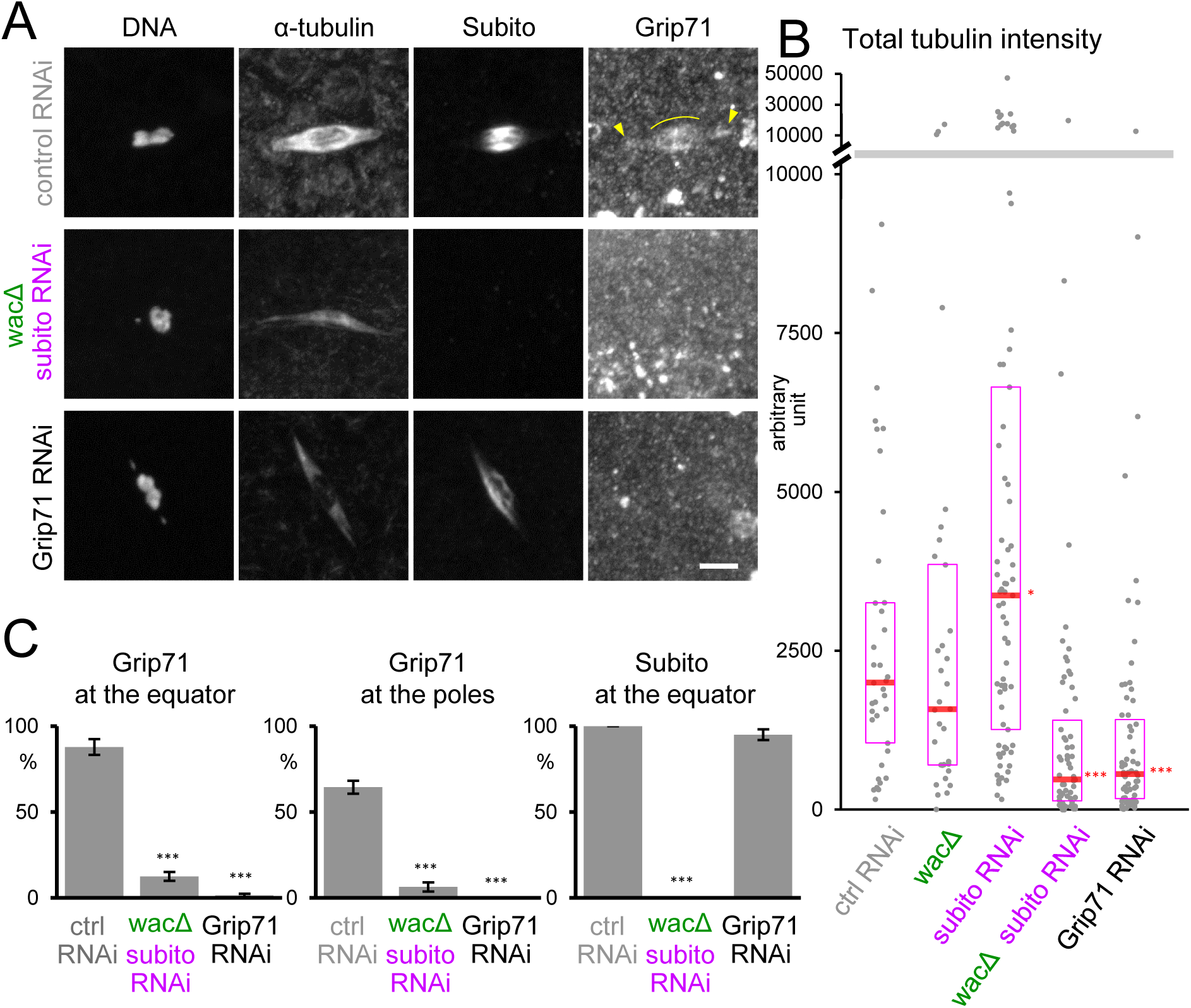
Subito and Augmin complement each other to generate microtubules. (A) Immunostaining of a meiotic spindle in mature oocytes from wild type expressing control (*white*) shRNA, a *wac* deletion mutant expressing *subito* shRNA, and wild type expressing *Grip71* shRNA. Arrowheads indicate Grip71 localisation at the spindle pole, and curved lines indicate Grip71 concentration on the spindle equator. Spindle microtubules were greatly reduced and Grip71 localisation on the spindle was lost in the *wac* deletion mutant expressing *subito* shRNA. These images were taken using the same settings and are shown without contrast enhancement. Bar=5 µm. (B) The total tubulin intensity of the meiotic spindle in each genotype (37, 33, 73, 63, 67 spindles). RNAi of the *white* gene was used as the control RNAi. The median is indicated by the central line, and the second and third quartiles are indicated by the box. The Y-axis is broken into two with different scales to show all data. Data were pooled from triplicate experiments. (C) The frequencies of meiotic spindles which have Grip71 concentration at the spindle equator and Grip71 localisation at at least one of the spindle poles, respectively, and Subito at the equator. The graphs show the means and standard errors from four repeats for each genotype (53, 89, 98 spindles in total). * and *** indicate significant differences from control RNAi (p<0.05, and 0.001, respectively).

### Grip71 mediates ectopic microtubule assembly induced by Subito lacking the N-terminal region

It was previously shown that over-expression of Subito lacking its N-terminal non-motor region in oocytes can induce ectopic spindles in the cytoplasm, away from the chromosomes (Jang et al., 2007; **Fig 4A**). Originally it was thought that mis-regulated Subito bundles cytoplamic microtubules, resulting in the formation of ectopic spindles (Jang et al., 2007). However, an alternative interpretation is that mis-regulated Subito ectopically recruits the γ-tubulin complex in the ooplasm and nucleates microtubules, independently of chromosomes. If this interpretation is correct, we predict that (1) Grip71 is recruited to the ectopic sites of microtubule assembly and that (2) the formation of the ectopic spindles depends on Grip71.

**Figure 4.**
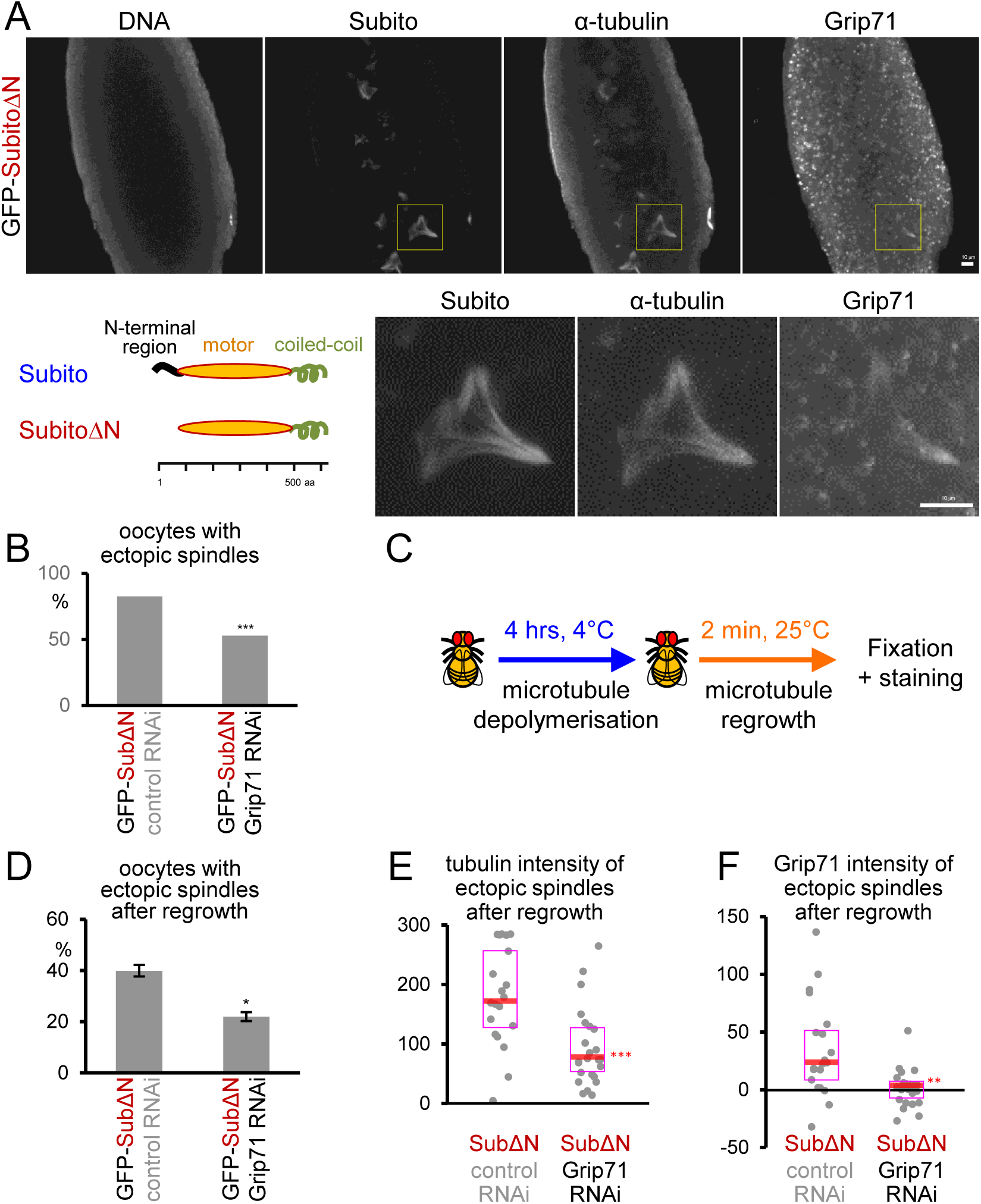
Grip71 mediates ectopic microtubule assembly induced by expression of Subito lacking the N-terminal region. (A) Grip71 is recruited to the ectopic spindles in oocytes expressing Subito lacking the N-terminal 89 residues (SubitoΔN). Immunostaining of mature wild-type oocytes expressing GFP-SubitoΔN that lacks the N-terminal non-motor domain as shown in the diagram. GFP-SubitoΔN was detected by an antibody against Subito. The boxed area is magnified below. Bars=10 µm. (B) The frequencies of oocytes which have ectopic spindles when GFP-SubitoΔN was expressed together with control (*white*) or Grip71 shRNA. 100 and 160 oocytes were examined, respectively. *** indicates a significant difference from wild type (Fischer’s exact test). (C) Schematic diagram of microtubule regrowth experiments. (D) The frequencies of oocytes with ectopic spindles when GFP-SubitoΔN was expressed together with shRNA against control (*white*) or *Grip71* after microtubule regrowth. The graph shows the means and standard errors from two repeats for each genotype (202 and 277 oocytes in total). * indicates a significant difference from wild type (p<0.05). (E,F) Signal intensities of α-tubulin and Grip71 on ectopic spindles above the cytoplasmic background after microtubule regrowth in each genotype (20, 24 oocytes). The graphs show the data from one of two repeated experiments, both of which showed decreases to similar levels in Grip71 RNAi oocytes. The median is indicated by the central line, and the second and third quartiles are indicated by the box. ** and *** indicates a significant difference from control RNAi (p<0.01 and 0.001).

To test whether Grip71 is recruited to ectopic spindles, we immunostained Grip71 in oocytes expressing Subito lacking the N-terminal region (SubitoΔN). As previously shown (Jang et al., 2007), SubitoΔN not only localised to the equatorial region of the meiotic spindle associated with chromosomes, but was also concentrated on ectopic spindles in oocytes. Importantly, Grip71 was recruited to ectopic spindles in addition to the equatorial region of the meiotic spindle (**Fig 4A**).

Next, to determine whether the formation of the ectopic spindles depends on Grip71, Grip71 was depleted by RNAi from oocytes expressing SubitoΔN (**Fig 4B**). Over 80% of oocytes expressing SubitoΔN and control shRNA form ectopic spindles. In contrast, when Grip71 is depleted, about a half of the oocytes expressing SubitoΔN failed to form ectopic spindles. To assess microtubule nucleation more accurately, microtubule regrowth experiments were carried out using cold treatment followed by warming (**Fig 4C**). Cold treatment abolished both chromosome-associated spindles and ectopic microtubules. Two minutes after warming, ∼40% of control oocytes displayed ectopic microtubule arrays associated with Grip71 foci, while fewer Grip71-depleted oocytes (∼20%) displayed ectopic arrays (**Fig 4D**). Furthermore, these fewer ectopic arrays formed in Grip71-depleted oocytes showed substantially lower tubulin intensity than the control (**Fig 4E**). Fewer and weaker microtubule arrays in Grip71-depleted oocytes indicates that ectopic microtubule assembly by SubitoΔN is largely dependent on Grip71. These results support our hypothesis that SubitoΔN induces ectopic microtubule nucleation through recruiting the γ-tubulin complex in the ooplasm.

### Subito and its associated proteins can induce microtubule nucleation *in vitro*

Our genetic and cytological studies support the hypothesis that Subito recruits γ-tubulin through Grip71 to nucleate spindle microtubules. To biochemically demonstrate that Subito and its associated proteins can nucleate microtubules, we set up the following *in vitro* assay using GFP-tagged Subito immunoprecipitated from ovaries incubated with pure pig brain α/β-tubulin dimer (**Fig 5A**).

**Figure 5.**
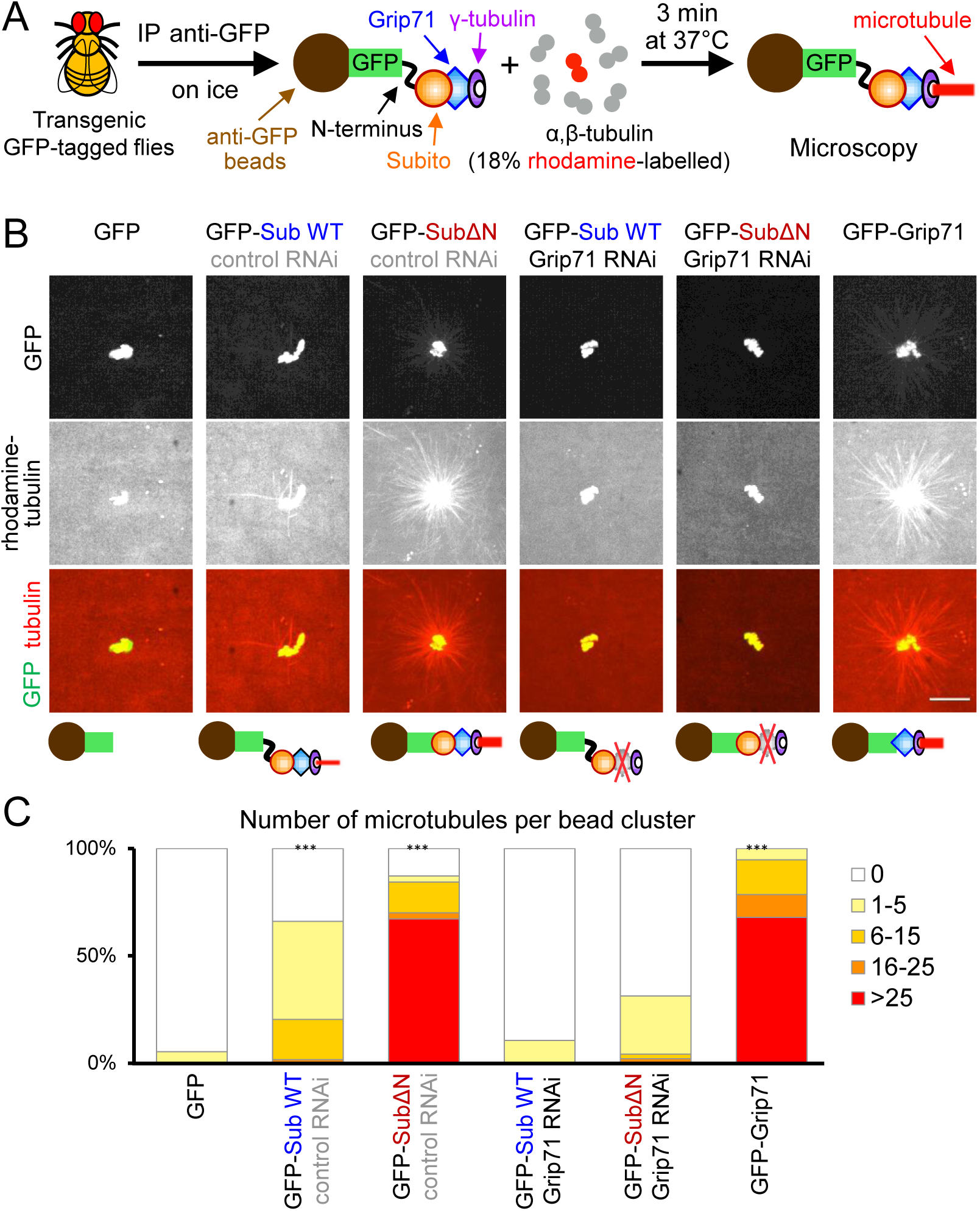
The Subito complex from ovaries can induce microtubule nucleation *in vitro*. (A) Schematic diagram of the assay of *in vitro* microtubule nucleation activity using an immunoprecipitated GFP-tagged protein (such as GFP-Subito) on beads incubated with pure pig α,β-tubulin dimer. (B) GFP-tagged protein was immunoprecipitated from oocytes expressing the protein alone or together with control (*white*) or *Grip71* shRNA. These images were taken using the same settings and the same alteration of the brightness and contrast was applied to all to show beads and microtubule asters clearly. Bar=10 µm. (C) The number of microtubules per bead cluster was counted for each genotype (75, 59, 70, 76, 48, 56 bead clusters). *** indicates a significant difference from GFP beads in the proportion of bead clusters with ≥6 microtubules. The graph shows the data from one of two repeated experiments, both of which showed similar results. In both experiments, GFP-SubitoΔN and GFP-Grip71 beads have a high microtubule nucleation activity, while GFP-Subito (GFP-Sub WT) beads have a lower nucleation activity. These activities depend on Grip71. GFP beads have virtually no nucleation activity.

Ovaries of flies expressing GFP-Subito (Subito WT) or GFP-SubitoΔN were used to immunoprecipitate each Subito variant using beads coupled with an anti-GFP antibody. In addition to GFP-tagged proteins, those oocytes also expressed shRNA against either *Grip71* or *white* (control). As negative and positive controls, we used GFP alone or GFP-Grip71 immunoprecipitated from corresponding ovaries. These beads were then incubated with pure α/β-tubulin dimer (18% of which were rhodamine-labelled) and GTP. After 3 minutes at 37°C, the reaction was fixed and observed under a fluorescence microscope.

As expected, the GFP-beads were associated with few microtubules, while the GFP-Grip71 beads were associated with an aster-like microtubule array (**Fig 5B,C**). Strikingly, GFP-SubitoΔN beads from control RNAi oocytes were associated with an aster-like microtubule array to a similar extent to GFP-Grip71 beads (**Fig 5B,C**). Interestingly, GFP-Subito WT beads from control RNAi oocytes were also associated with some microtubules but to a much lesser extent than GFP-SubitoΔN beads (**Fig 5B,C**). This stronger *in vitro* nucleation activity of GFP-SubitoΔN is consistent with its ability to induce ectopic microtubules *in vivo*.

It is formally possible that microtubules around the GFP-SubitoΔN beads were spontaneously nucleated in the solution and then captured by the beads, rather than nucleated by the beads. To exclude this possibility, we counted the number of free microtubules not associated with beads (**Fig S2**). If microtubules were spontaneously nucleated and then captured, fewer free microtubules should be observed in the experiments using GFP-SubitoΔN beads than with GFP-beads. In contrast, if GFP-SubitoΔN beads nucleated microtubules (some of which were detached), more free microtubules should be observed than GFP-beads. Indeed, we observed many more free microtubules in the experiment using GFP-SubitoΔN beads than using GFP-beads (**Fig S2**). This confirmed that GFP-SubitoΔN beads induced microtubule nucleation, rather than simply capturing spontaneously nucleated microtubules.

To further confirm that these microtubule asters formed around the GFP-SubitoΔN or GFP-Subito WT beads were due to microtubule nucleation, we tested whether the γ-tubulin subunit Grip71 is required for aster formation. GFP-SubitoΔN or GFP-Subito WT was immunoprecipitated from ovaries simultaneously expressing GFP-SubitoΔN and shRNA against Grip71. None or a few microtubules were assembled around the GFP-SubitoΔN or GFP-Subito WT beads from Grip71-depleted oocytes (**Fig 5B,C**). These results demonstrate that microtubule assembly mediated by the GFP-SubitoΔN or GFP-Subito WT beads depends on Grip71. Therefore, we conclude that Subito and its associated proteins can nucleate microtubules from pure α/β-tubulin *in vitro*, and that this nucleation activity is inhibited by the N-terminal region of Subito.

### Subito interaction with the γ-tubulin complex is suppressed by its N-terminal region

Genetic and cytological analysis in this study showed that the Subito N-terminal region suppresses the Subito activity and prevents ectopic microtubule nucleation in oocytes. Furthermore, the N-terminal region suppresses *in vitro* microtubule nucleation activity of Subito. To gain a mechanistic insight, we tested whether the interaction between Subito and the γ-tubulin complex is regulated by the N-terminal region. Proteins co-immunoprecipitated with GFP-Subito or GFP-SubitoΔN were analysed by immunoblot using Grip71 and γ-tubulin antibodies (**Fig 6**).

**Figure 6.**
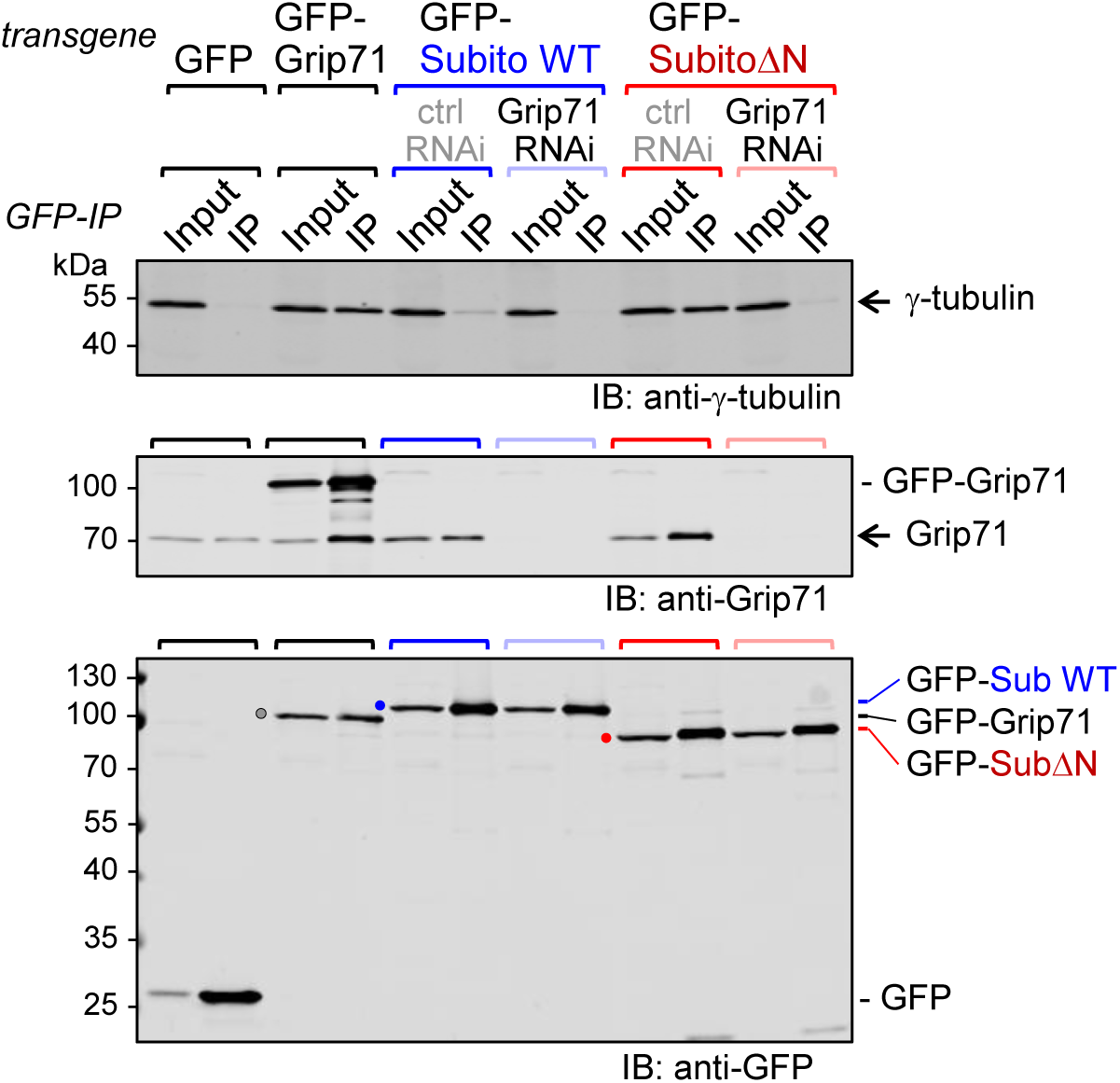
Subito interaction with Grip71 is negatively regulated by its N-terminal region. GFP-tagged protein was immunoprecipitated from oocytes expressing the protein alone or together with control (*white*) or *Grip71* shRNA in the presence of phosphatase inhibitors, followed by immunoblots (IB). 2.5% of Input relative to the immunoprecipiated fraction (IP) was loaded. γ-tubulin and Grip71 were efficiently co-immunoprecipiated with GFP-SubitoΔN, but only weakly with GFP-Subito. Co-immunoprecipitation of γ-tubulin depends on Grip71. This figure shows the immunoblot from one of two repeated experiments, both of which provided similar results.

Strikingly, we found that Grip71 and γ-tubulin were co-immunoprecipitated with GFP-SubitoΔN much more efficiently than with GFP-Subito (**Fig 6**). This demonstrated that the N-terminal region suppresses the interaction between Subito and the γ-tubulin complex, which explains why SubitoΔN is hyperactive in nucleating microtubules *in vivo* and *in vitro*. When Grip71 was depleted from ovaries, γ-tubulin was not co-immunoprecipitated with either GFP-SubitoΔN or Subito, confirming that Grip71 mediates the interaction between Subito and the γ-tubulin complex (**Fig 6**). These results show that the N-terminal region inhibits the interaction between Subito and Grip71, hence between Subito and the γ-tubulin complex.

## Discussion

Our *in vivo* and *in vitro* analysis uncovered a novel microtubule nucleation pathway specific to oocytes, which is mediated by the conserved kinesin-6 Subito/MKlp2. Subito recruits the γ-tubulin complex to the spindle equator via Grip71/NEDD1, acting complementarily to Augmin which fulfils this role at the spindle poles. A loss of either pathway does not reduce bulk spindle microtubules, but together they are essential for the assembly of most spindle microtubules in oocytes. We demonstrated that Subito can interact with the γ-tubulin complex through Grip71 and induce microtubule nucleation from pure tubulin dimers *in vitro*. Interestingly the N-terminal region of Subito suppresses its interaction with the γ-tubulin complex, and prevents ectopic assembly of microtubules in oocytes. Therefore, this novel microtubule nucleation pathway is central to assemble the meiotic spindle around chromosomes and, as importantly, to suppress ectopic microtubule nucleation in the ooplasm.

We propose the following model of spatially restricted spindle assembly in *Drosophila* oocytes (**Fig 7**). In the vicinity of chromosomes, Subito binds to microtubules in the spindle equator and recruits the γ-tubulin complex through Grip71 to nucleate new microtubules. In the same manner, Augmin localises to the spindle poles where it nucleates microtubules. Away from chromosomes, the N-terminal region of Subito suppresses its interaction with the γ-tubulin complex and therefore prevents ectopic microtubule nucleation.

**Figure 7.**
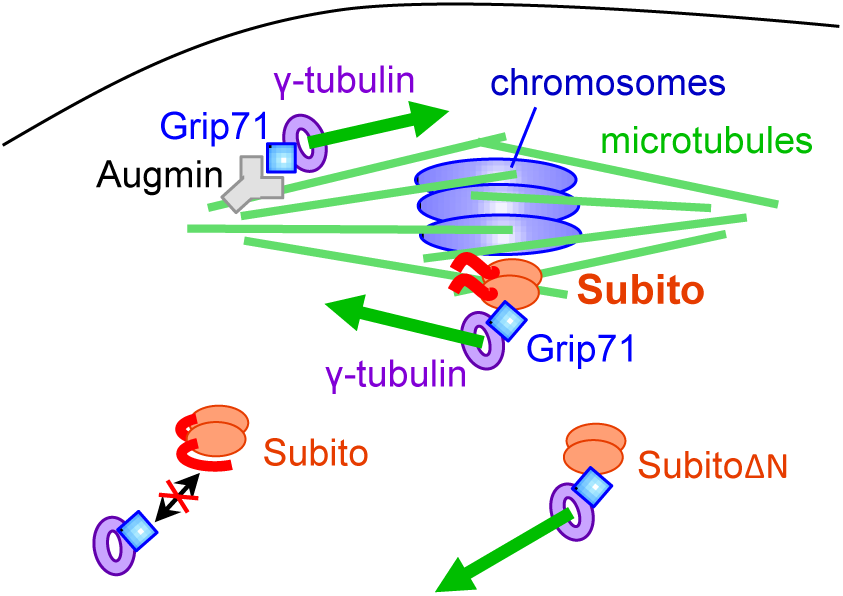
Subito recruits the γ-tubulin complex to nucleate microtubules in the spindle equator in oocytes. Schematic model of meiotic spindle assembly in oocytes. Subito and Augmin recruit the γ-tubulin complex to the spindle equator and poles, respectively, through Grip71. This nucleates a new microtubule to amplify spindle microtubules. These two pathways work complementarily to assemble most of the spindle microtubules in oocytes. The N-terminal region of Subito suppresses the interaction with the γ-tubulin complex away from chromosomes, but this suppression is removed near meiotic chromosomes. Subito lacking the N-terminal region constitutively associates with the γ-tubulin complex and induces ectopic microtubule nucleation away from chromosomes in the ooplasm.

Taking advantage of *Drosophila* oocytes, we combined genetic, cytological and biochemical approaches to uncover a novel oocyte-specific microtubule nucleation pathway. Unique features of oocytes, such as lack of centrosomes and a large volume, are also characteristics of oocytes from other species including humans. Furthermore, all proteins mentioned here, such as Subito/MKlp2, Grip71/NEDD1, Augmin and the γ-tubulin complex, are widely conserved (Jang et al., 2005; Haren et al., 2006; Lüders et al., 2006; Uehara et al., 2009; Lawo et al., 2009; Kollman et al., 2011; Teixidó-Travesa et al., 2012). Therefore, a similar mechanism to this novel nucleation pathway is likely to be operating in oocytes of other species to assemble the meiotic acentrosomal spindle.

Subito provides the first example of a kinesin to recruit a microtubule nucleator. As a kinesin, Subito also binds to microtubules. Thus, by combining microtubule binding and nucleation activities, it could locally increase the number of microtubules in an exponential manner to trigger rapid assembly of a very dense microtubule network (which is called microtubule amplification; Zhu et al., 2008; Goshima et al., 2010). Augmin was the only previously known factor which mediates microtubule amplification for spindle assembly (Meunier and Vernos, 2016; Prosser and Pelletier, 2017). In oocytes, we showed that the Subito pathway acts complementarily to Augmin pathway to assemble most of the spindle microtubules in oocytes. In mitosis, however, Subito microtubule nucleation pathway seems to be inactive or negligible, as very few microtubules are assembled in the absence of both centrosomes and Augmin (Goshima et al., 2008; Meireles et al., 2009; Wainman et al., 2009). Moreover, Subito depletion does not result in significant spindle defect in mitotic cells lacking centrosomes (Moutinho-Pereira et al., 2013). Therefore, this new role of Subito in microtubule nucleation appears to be specific to oocytes.

During animal mitosis, centrosomes are major microtubule nucleating sites (Prosser and Pelletier, 2017) and Augmin nucleates randomly along the spindle (Goshima et al., 2008). Based on our findings, we hypothesise that the absence of the centrosomal activity in oocytes is compensated by two oocyte-specific adaptations of Augmin and Subito. Firstly, oocyte-specific localisation of Augmin at the spindle poles compensates for the absence of nucleation activity of centrosomes from the poles (Meireles et al., 2009; Colombie et al., 2013). Secondly, oocyte-specific Subito pathway compensates for the absence of microtubule nucleation by Augmin in the equator region.

Once microtubules are nucleated in the spindle equator, microtubules are cross-linked and their polarity is sorted out by microtubule motors (Bennabi et al., 2016). As the mammalian orthologue of Subito, MKlp2, has been shown to bundle microtubules (Neef et al., 2003), Subito can potentially coordinate nucleation, cross-linking and polarity sorting during meiotic spindle assembly.

In addition, our study highlights a central role of Subito in spatially restricting microtubule amplification to the vicinity of chromosomes in *Drosophila* oocytes. This spatial regulation is crucial, as oocytes commonly contain exceptionally large cytoplasmic volumes. We showed that the N-terminal non-motor region of Subito suppresses ectopic microtubule nucleation by inhibiting the interaction with Grip71, the subunit responsible for recruiting the γ-tubulin complex. It is possible that the N-terminal region may recruit a protein which inhibits the interaction with Grip71. However, there are many examples of auto-inhibition of kinesins by their non-motor region (Verhey and Hammond, 2009). It is therefore likely that a signal from chromosomes locally abolishes auto-inhibition of the Subito nucleation activity in order to trigger spindle microtubule nucleation in a spatially confined manner. In simple terms, chromosomes switch on Subito-dependent nucleation, while this pathway remains switched off in the cytoplasm of the oocyte. Ran-GTP or Aurora B activity could potentially act as the signal from the meiotic chromosomes to remove auto-inhibition of Subito (Bennabi et al., 2016; Radford et al., 2017). Taken together, tight regulation of Subito appears critical to assemble spindle microtubules around the chromosomes and to prevent their assembly anywhere else.

## Methods

### Recombinant DNA techniques

Standard molecular techniques (Sambrook et al., 1989) were followed throughout. cDNA of each gene of interest was cloned firstly into pENTR/D-TOPO using pENTR Directional Topo Cloning Kit (Invitrogen). The absence of unwanted mutations was confirmed by DNA sequencing using BigDye (PerkinElmer). cDNAs were then recombined into Gateway destination vectors using LR Clonase II reaction following the manufacturer’s protocol (Invitrogen). To generate transgenic flies expressing a GFP-tagged protein under the UASp promoter, we used φPGW modified from destination vector pPGW of the Murphy’s Gateway collection (https://emb.carnegiescience.edu/drosophila-gateway-vector-collection) by adding the φC31 attB recombination site (Beaven et al., 2017).

To generate the *Grip71* entry clone, an entire coding region was amplified from cDNA (RE05579) by PCR. This cDNA contains a cytosine deletion at 2,134 bp that induces a premature stop codon. The cytosine has been added back to restore Grip71 complete coding region (1-646 amino acids followed by a stop codon) using the Quick Change II XL Site-Directed Mutagenesis kit (Promega) and the following primers: CCAACATTTCCGTGGCCAGCAGCACAGGAGGCGGCAGCG and CGCTGCCGCCTCCTGTGCTGCTGGCCACGGAAATGTTGG.

To generate the *subito* entry clone, a full-length coding region (encoding 1-628 amino acids followed by a stop codon) or a part of the coding region (SubitoΔN encoding 90-628 amino acids followed by a stop codon) was amplified from cDNA (LD35138) by PCR.

For flies expressing *Grip71* shRNA, a Walium22 plasmid carrying shRNA against position 63 of the 5’UTR of the *Grip71* gene was generated following TRiP protocol (http://hwpi.harvard.edu/files/fly/files/2ndgenprotocol.pdf?m=1465918000) using the following primers: CTAGCAGTGTGTAAATATCTGAAGAAATATAGTTATATTCAAGCATATATTTCTTCAGATATTTACACGCG and AATTCGCGTGTAAATATCTGAAGAAATATATGCTTGAATATAACTATATTTCTTCAGATATTTACACACTG.

For Figure 4B, an additional line expressing a different shRNA directed against position 375 of the coding region of *Grip71* gene was generated using the following primers: CTAGCAGTGAGCGGTTGTGTTAAGCTATATAGTTATATTCAAGCATATATAGCTTAACACAACCGCTCGCG and AATTCGCGAGCGGTTGTGTTAAGCTATATATGCTTGAATATAACTATATAGCTTAACACAACCGCTCACTG.

For RNAi in S2 cells, dsRNA against the β-lactamase gene was used as a control as previously described (Syred et al., 2013). For *Grip71* RNAi in S2 cells (Fig S1A, B), double-stranded RNAs were amplified in the first PCR using the following pairs of primers containing half of the T7 promoter and the full T7 promoter sequence was added in the second PCR using the primer, CGACTCACTATAGGGAGAGCGGGGGATTCTCTTTGT. In the first PCR, the region 504-925 bp of Grip71 coding sequence was amplified using CGACTCACTATAGGGAGACCGAGCAAACGCTTTCAT and CGACTCACTATAGGGAGATGCTGGTCATCCCCACTT for Grip71 RNAi 1, and the region 1062-1479 bp of Grip71 coding sequence was amplified by CGACTCACTATAGGGAGATTGGTTACGGGGTGTCAA and GAATTAATACGACTCACTATAGGGAGA for Grip71 RNAi 2. The PCR products were *in vitro* transcribed and purified using MEGAscript T7 High Yield Transcription Kit (Ambion) following the manufacturer’s instructions.

### *Drosophila* techniques

Standard methods of fly handling (Ashburner et al., 2005) were used. *w^1118^* was used as wild type in this study. For a *Grip71* deletion mutant, we used *Dgp71WD^120^* over a deficiency from a cross between *Dgp71WD^120^/SM6A* (Reschen et al., 2012) and a deficiency *Df(2L)ED1196/SM6A*. The *wac* deletion mutant line used in this study was a homozygote from a *wac^Δ12^/TM6C* stock (Meireles et al., 2009). A line expressing shRNA against *subito* (GL00583) was used to inhibit *subito* expression. Two lines expressing shRNA against *Grip71* 5’UTR or the coding region (this study) were used to inhibit *Grip71* expression. In Figure 4B, as both lines gave very similar results, they were pooled together for the analysis. For other experiments, the line expressing shRNA against *Grip71* 5’UTR was used. The line (GL00094) expressing shRNA against *white* was used as a negative control (control RNAi) in this study. shRNA and transgenes encoding for GFP-Grip71, GFP-Subito WT and GFP-SubitoΔN under the control of the UASp promoter were expressed using one copy of a female germline specific GAL4 driver, *nos-GAL4-VP16 MVD1*, except Figure 2. In Figure 2, a maternal-tubulin GAL4 driver V37 was used in one of the three experiments, but as it gave a similar result to the other two using MVD1, all three experiments were analysed together. To express GFP-Subito WT or GFP-SubitoΔN together with *white* shRNA (a negative control) or *Grip71* shRNA, a recombinant chromosome between two transgenes (both on the third chromosome) was generated and then crossed with MVD1. Recombinant chromosomes were established by selecting and confirming using visible markers and PCR after meiotic recombination in females.

For generating transgenic fly lines, φPGW carrying cDNA encoding the full-length Subito (Subito WT) and SubitoΔN, and an empty φPGW, were integrated at the landing site VK33 (Venken et al., 2006) on the third chromosome using φC31 integrase by Best Genes Inc.. Similarly, φPGW carrying cDNA encoding the full-length Grip71 and Walium22 carrying shRNA targeting 5’ UTR or the coding sequence of Grip71 were integrated at attP2 (Groth et al., 2004) on the third chromosome.

### Immunological reagents and techniques

For generating an anti-Grip71 antibody, an entry vector (pENTR/D-TOPO) carrying the Grip71 full coding sequence from a cDNA (RE05579) was used. The cDNA has a mutation that leads to a premature stop codon after 493rd amino acid of Grip71. For expression of GST-tagged or MBP-tagged truncated Grip71 (Grip71*) in bacteria, the entry plasmid was recombined with expression vectors (pGEX4T1 and pMALc2) adapted to Gateway via LR Clonase (Invitrogen), respectively. *BL21(DE3) pLysS* containing GEX4T1-Grip71* were cultured overnight at 18°C in the presence of 1 mM IPTG. Bacteria were lysed by sonication in PBS + 0.5% Triton X-100 and spun down at 4,000 rpm for 5 min. The pellet contained most of GST-Grip71* which were largely insoluble in this condition. After a wash in the same buffer, the pellet was run on an SDS protein gel, and eluted from the gel by diffusion in 0.2M NaHCO_3_ + 0.02% SDS. A total of 2.5 mg of protein (4 injections) was used to immunise a rabbit by Scottish National Blood Transfusion Service. The final anti-serum was then affinity-purified using MBP-Grip71* on nitrocellulose membranes as previously described (Smith and Fisher, 1984; Clohisey et al., 2014).

The following primary antibodies were used for immunofluorescence microscopy (IF) and immunoblots (IB) in the study: anti-GFP (rabbit polyclonal A11122; Thermo Fisher Scientific; 1:250 for IF and IB), anti-Subito (rat polyclonal against GST-Subito full-length; Loh et al., 2012; 1:250), anti-Grip71 (rabbit polyclonal affinity purified; this study; 1:2-1:10 for IF and 1:50 for IB), anti-γ-tubulin (mouse monoclonal GTU88; Sigma-Aldrich; 1:250 for IB), anti-α-tubulin (mouse monoclonal DM1A; Sigma-Aldrich; 1:250 for IF). Alexa 488-, Alexa 647, Cy3-, Cy5-conjugated secondary antibodies were used (1:250 to 1:1000; Jackson Laboratory or Molecular Probes) for IF, and IRDye 800CW conjugated goat anti-rabbit (1:20,000), IRDye 680LT conjugated goat anti-rabbit (1:15,000), and IRDye 800CW conjugated goat anti-mouse (1:20,000) secondary antibodies (LI-COR) were used for IB.

For immunoblotting, SDS-PAGE (sodium dodecyl sulfate polyacrylamide gel electrophoresis) were carried out and proteins on the gel were transferred onto a nitrocellulose membrane following a standard procedure (Sambrook et al., 1989). Total proteins on the membrane were stained with MemCode reversible staining kit (Thermo-Fisher) or Ponceau (Sigma-Aldrich). After being destained, the membrane was incubated with primary antibodies followed by fluorescent secondary antibodies (LI-COR). The signals were detected with an Odyssey imaging system for Figure 2B, S1B,E and an Odyssey CLx imaging scanner (LI-COR) for Figure 6. The brightness and contrast were adjusted uniformly across the entire area without removing or altering features.

For immunoblotting of ovaries, 10 ovaries or 100 oocytes were dissected from mature females in methanol. Methanol was replaced by 50 μl of water and 50 μl of boiling 2x sample buffer (50 mM Tris-HCl pH6.8; 2% SDS; 10% glycerol; 0.1% Bromophenol Blue; 715 mM 2-mercaptoethanol). Ovaries were homogenised in a microtube using a pestle (Eppendorf). The equivalent of 20 oocytes (20 μl, Figure S1E) or 1 ovary (10 μl, Figure 2D) were loaded on a gel for SDS-PAGE followed by immunoblot. For S2 cells, 5 million cells were spun down at 500xg for 10 minutes, washed once with PBS at room temperature then resuspended in 50 μl of water and 50 μl of boiling 2x sample buffer. The equivalent of 500,000 cells were loaded on a gel for SDS-PAGE followed by immunoblot (Figure S1B).

### Cytology and image analysis

For immunostaining of mature oocytes, freshly eclosed females were kept with males for 3-5 days at 25°C with an excess of dried yeast. Ovaries from matured females were dissected in methanol and, after removing the chorion by sonication, oocytes were stained as previously described (Cullen and Ohkura, 2001). Under this experimental condition, most oocytes were arrested at stage 14.

Microtubule regrowth experiments in Figure 4C-F were carried out as follows. 24 microtubes containing 1-2 mature females each were incubated on ice for 4 hours to depolymerise microtubules. The absence of microtubules in oocytes after 4 hours on ice was confirmed by immunostaining. Each tube was transferred one-by-one to water at 25°C, and after 2 minutes, ovaries were dissected in methanol and processed for immunostaining.

Immunostained oocytes were imaged with a confocal scan head, LSM510Exciter (Zeiss; Figure 1A, 2A,C, 4A, S1C) or LSM800 with GaAsP photomultipliers (Zeiss; Figure 3, 4B,D,E,F), attached to an Axiovert 200M (Zeiss) using 63x/NA1.40 Plan-ApoChromat objective (oil) and 0.5 μm Z-step acquisition. The capturing resolution was set for 100 nm per pixel. Representative images are shown in figures after maximum intensity projection of multiple Z-planes including the entire spindle. The contrast and brightness of microscope images shown in the figures were unaltered. When the signal intensity is compared (Figure 3A,B), the images were taken using the same settings.

For *Drosophila* S2 cells (Schneider, 1972), about 1 million cells were aliquoted into each well of a 6-well petri dish with 1 ml of Serum Free Medium (Schneider). 15 μg of dsRNA is added to each well. After 1 hour, 2 ml of the medium containing 10% FCS was added to the cells. Three days later the cells were transferred to concanavalin A coated coverslips (18×18 mm, VWR), fixed with a cold solution containing 90% methanol, 2.88% paraformaldehyde and 5mM NaHCO_3_ (pH9), and stained as previously described (Dzhindzhev et al., 2005). A single Z section of S2 cells was imaged with a CCD camera (Orca; Hamamatsu) attached to an upright microscope (Axioplan2) controlled by OpenLab (Improvision) using a 100x/NA1.4 Plan-ApoChromat objective.

For Figure 3, the total tubulin intensity was measured on spindles imaged with the same setting (A). If some signals were saturated, a second image was taken with a lower laser intensity (B). To compensate for the lower laser power, a laser coefficient was calculated by the ratio (A/B) of the total signal intensity of three spindles without saturated signals taken by these two settings. Tubulin signal intensity was measured from maximum intensity projection of a Z-stack. A region S mainly containing the spindle was drawn, and the total intensity (IntDenS) and the area size (AreaS) was measured. A second region L containing the spindle and the surrounding background was drawn, and the total tubulin intensity (IntDenL) and the area size (AreaL) were measured. The following formula was then applied to calculate the total tubulin intensity for each spindle: IntDenS-AreaS*(IntDenL-IntDenS)/(AreaL-AreaS). For the overexposed spindles, the tubulin intensity was measured from an image captured using setting B and the following formula was applied: LaserCoefficient*[IntDenS-AreaS*(IntDenL-IntDenS)/(AreaL-areaS)]. Two-tailed Wilcoxon rank-sum test was used for testing statistical significance.

For Figure 4E,F, S1D, the tubulin and Grip71 signal intensities were measured from the maximum intensity projection of a Z-stack as follows. A small region of interest (region C) was drawn on the ectopic spindle (Figure 4E,F) or near the chromosomes on the spindle (Figure S1D), and the total signal intensities of tubulin and Grip71 were measured. A region of interest with the identical shape and size (region D) was drawn in a background region and the total signal intensities were measured. The specific signal intensity was calculated by subtracting the total intensity of region D from the total intensity of region C. For Figure S1D, the specific signal intensity of Grip71 was then normalised by dividing it by the specific tubulin signal intensity of the same spindle. Two-tailed Wilcoxon rank-sum test was used for testing statistical significance.

For Figure 1B, 2E and 3C, accumulation of Grip71 and Subito in the spindle equator and poles was judged in comparison with the intensity distribution of tubulin signals by visual inspection of each Z-plane. To check the validity of classification, a series of raw images on which Figure 1B is based were independently examined by another researcher in a blind manner, which gave a similar result. Multiple independent batches of immunostaining were carried out for each genotype, and the mean and standard error (SEM) were calculated for the percentage of spindles which showed the accumulation in each replicate. Two-tailed unpaired t-test was used for testing statistical significance.

For Figure 4B,D, the entire oocytes were observed for the presence or absence of dense microtubule arrays not associated with the meiotic chromosomes (“ectopic spindles”). Two independent batches of immunostaining were carried out for each genotype in D, and the mean and standard error (SEM) were calculated for the percentage of oocytes which have visible ectopic spindles in each replicate. Two-tailed unpaired t-test was used for testing statistical significance. As only one batch of immunostaining was carried out for GFP-SubitoΔN + *white* shRNA in B, statistical significance was tested by Fisher’s exact test.

### Immunoprecipitation and microtubule nucleation assay

Immunoprecipitation and microtubule nucleation assay shown in Figure 5, 6 and S2 were carried out as follows. Flies expressing GFP, GFP-Grip71, GFP-Subito WT + *white* shRNA, GFP-Subito WT + *Grip71* shRNA, GFP-SubitoΔN + *white* shRNA, or GFP-SubitoΔN + *Grip71* shRNA were matured with dried yeast for 3-5 days at 25°C. 20 pairs of ovaries from each genotype were dissected in PBS + 2 mM EGTA and frozen with a minimum carry-over of the buffer on the side of a microtube prechilled on dry ice. Microtubes were snap frozen in liquid nitrogen and stored at −80°C.

For immunoprecipitation, 20 pairs of frozen ovaries were resuspended in 400 μl of lysis buffer (20 mM Tris-HCl pH7.5, 50 mM NaCl, 5 mM EGTA, 1 mM DTT supplemented with 1 mM PMSF, inhibitor of protease (Roche, 1 tablet for 10 ml)) + 1 μM okadaic acid, 10 mM p-nitrophenyl phosphate and 0.5% Triton X-100 and homogenised on ice with a Dounce tissue grinder (1 ml). Lysates were incubated on ice for 30 minutes and transferred in a microtube before being centrifuged for 30 minutes, at 13,000 rpm at 4°C. 10 μl was kept as Input and mixed with 10 μl of boiling 2x sample buffer, and the rest was used for the following immunoprecipitation. 11 μl of anti-GFP magnetic beads (Chromotek) were washed three times with cold lysis buffer + 0.1% Triton X-100 before being mixed with lysate supernatant for 1 hour at 4°C on a rotating wheel. The beads were washed once with cold lysis buffer containing 0.1% Triton X-100, 1 μM okadaic acid and 10 mM p-nitrophenyl phosphate. The beads were additionally washed twice with cold lysis buffer + 0.1% Triton X-100 and twice with cold BRB80 (80 mM PIPES, 1 mM MgCl_2_, 1 mM EGTA, pH 6.8). The beads were resuspended in 10 μl of cold BRB80 and kept on ice for the following microtubule nucleation assay.

For nucleation assay, pig unlabelled pure tubulin (Cytoskeleton T240-B) and pig rhodamine-labelled pure tubulin (Cytoskeleton TL590M-A) were resupended in BRB80 + 1 mM GTP to a final concentration of 50 μM and 100 μM, respectively. Unlabelled and labelled tubulin were mixed (4.5:1 in molar concentration) to a final concentration of 55 μM then diluted in BRB80 + 1 mM GTP to a final concentration of 36.6 μM. 9 μl of this mix was added to 1 μl of the beads for a final concentration of 33 μM of tubulin. The beads and tubulin were incubated for 3 minutes at 37°C to allow polymerisation. To stop the reaction, 10 μl of BRB80 at 37°C containing 1 mM GTP and 2% glutaraldehyde (Sigma-Aldrich G6403) was added to 10 μl of the reaction and gently mixed using a pipet tip, the end of which has been cut off. 10 μl of the reaction was mixed with 10 μl of BRB80 + 30% glycerol (15% glycerol final) to increase the viscosity and reduce movement of the beads. 4 μl of the reaction was mounted between two coverslips (24×50mm and 18×18mm, VWR).

Images were taken using a PlanApochromat lens (63x/1.4NA; Zeiss) attached to an Axiovert 200M microscope (Zeiss) with a spinning disc confocal scan head (CSU-X1; Yokogawa) operated by Volocity (PerkinElmer). 20-40 random fields were imaged with 488 nm laser (20%) to detect the GFP beads and 561 nm laser (20%) to detect the rhodamine labelled microtubules (250 ms exposure, 0.8 μm Z-interval). Microtubules associated with each bead cluster were counted from the maximum intensity projection of Z-stack. The numbers of free microtubules were counted in the single Z-plane where the GFP intensity was the highest (Figure S2). Two-tailed unpaired Wilcoxon rank-sum test was used for testing statistical significance. The images were taken using the identical setting and are shown in Figure 5B after applying the identical contrast and brightness enhancement. After the experiment, about 10 μl of the remaining beads were mixed with 10 μl of boiling 2x sample buffer. 10 μl of Input and beads (IP) were loaded on a gel for SDS-PAGE followed by immunoblot (Figure 6).

Immunoprecipitation shown in Figure 2B was carried out as follows. Ovaries were dissected from matured females for each condition in PBS + 0.5% Triton X-100. 25 pairs of ovaries were resuspended in 500 μl of lysis buffer. Ovaries were homogenised on ice using a Dounce tissue grinder (1 ml). Lysates were incubated on ice for 30 minutes and transferred in a microtube before being centrifuged at 13,000 rpm for 30 minutes at 4°C. 50 μl of the supernatant was kept as Input and mixed with 50 μl of boiling 2x sample buffer, and the rest is kept for the following immunoprecipitation. 30 μl of anti-GFP magnetic beads were washed three times with cold lysis buffer + 0.1% Triton X-100 before being mixed with the lysate supernatant for 2 hours at 4°C on a rotating wheel. The beads were isolated with a magnetic rack and washed three times with cold lysis buffer + 0.1% Triton X-100. The beads were resuspended in 50 μl of cold lysis buffer + 0.1% Triton X-100 and mixed with 50 μl of 2x boiling sample buffer. 10 μl of the Input and beads (IP) were loaded on a gel for SDS-PAGE followed by immunoblot.

## Data availability

The datasets generated and/or analysed during the current study are available from the corresponding author upon request.

## Acknowledgements

We are grateful to all the members of the Ohkura lab and D. Finnegan for their help and useful feedback, to E. Peat for her technical help and to J. Raff, the Bloomington Drosophila Stock Center and Resource Center (NIH P40OD018537, 2P40OD010949-10A1) and the Transgenic RNAi Project at Harvard Medical School (NIH/NIGMS R01-GM084947) for fly stocks and reagents. The work is supported by Wellcome Trust Senior Research Fellowships (081849, 098030) and Investigator Award (206315) to HO. The Wellcome Centre for Cell Biology is supported by a core grant (092076, 203149).

## Author contributions

PR designed and performed experiments, analysed the data and wrote the manuscript; HO designed experiments, analysed the data, and wrote the manuscript.

## Supplementary information

**Figure S1.**
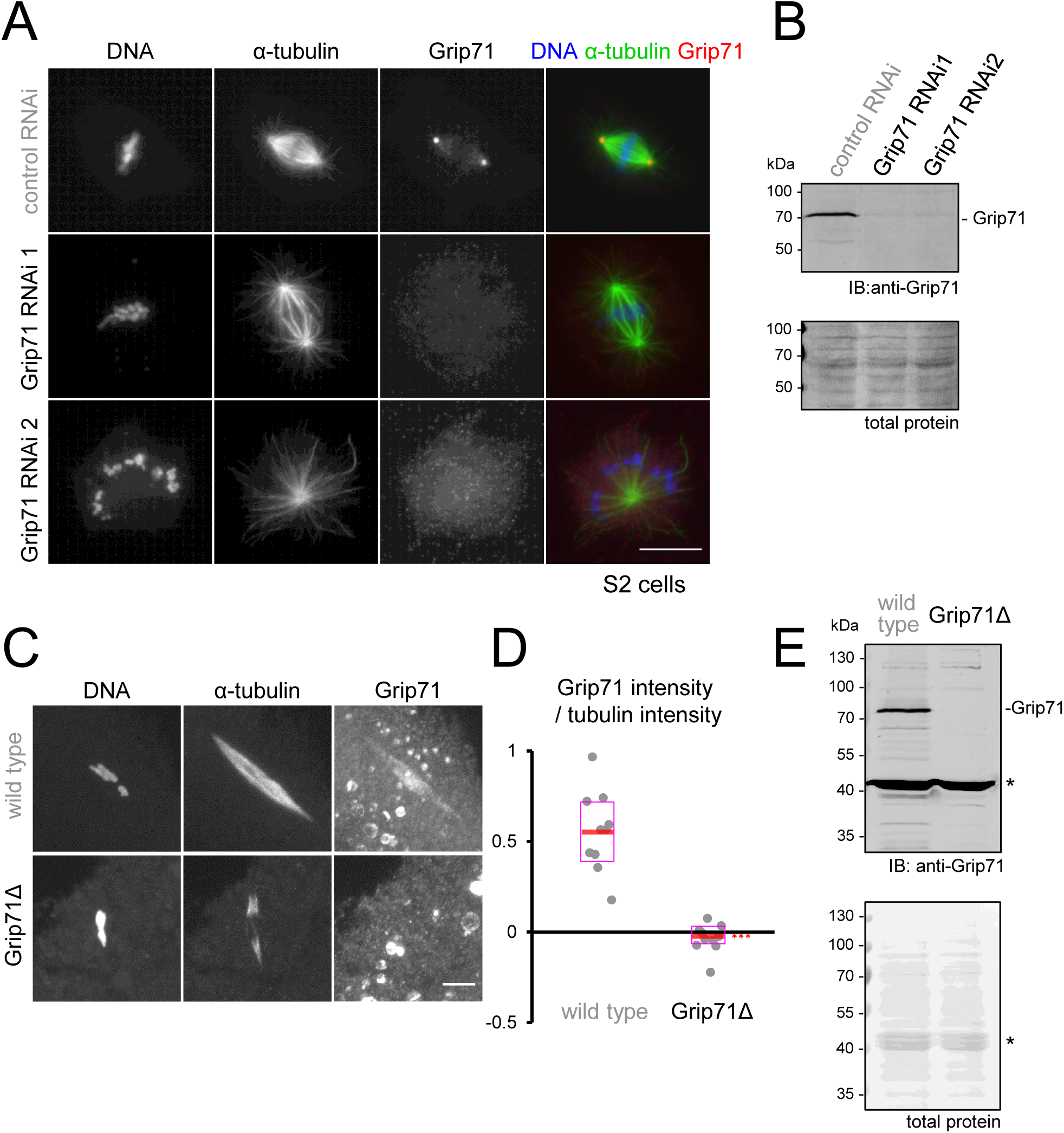
Specificity of the anti-Grip71 antibody. (A) Immunostaining of S2 cells incubated with control dsRNA (against β-lactamase gene) and two non-overlapping dsRNA against *Grip71*. The rabbit antibody against Grip71 stained spindle poles in control RNAi, but the signal greatly reduced in Grip71 RNAi cells. Bar=10 µm. (B) Immunoblot of S2 cells after RNAi using the anti-Grip71 antibody. The anti-Grip71 antibody gave a band of the expected size, the intensity of which was greatly reduced following Grip71 RNAi. (C) Immunostaining of a meiotic spindle in mature oocytes from wild type and a Grip71 deletion mutant using the anti-Grip71 antibody. Bar=5 µm. (D) Grip71 signal intensity relative to α-tubulin signal intensity near the chromosomes on the spindle in mature wild-type and *Grip71Δ* oocytes (10 spindles each). The signal intensity on the spindle in each oocyte was normalised using the background (cytoplasmic) intensity. The median is indicated by the central line, and the second and third quartiles are indicated by the box. The graph shows the data from one of two repeated experiments, both of which showed similar decreases in *Grip71Δ* oocytes. *** indicates a significant difference from wild type (p<0.001). (E) Immunoblot of ovaries from wild type and Grip71 deletion mutant probed by the anti-Grip71 antibody. The 75 kDa band disappeared in the deletion mutant and roughly coincides with the predicted molecular weight of Grip71, indicating that this band represents Grip71. On the other hand, the 42 kDa band (*) that coincides with yolk proteins was unchanged, indicating that this band is non-specific.

**Figure S2.**
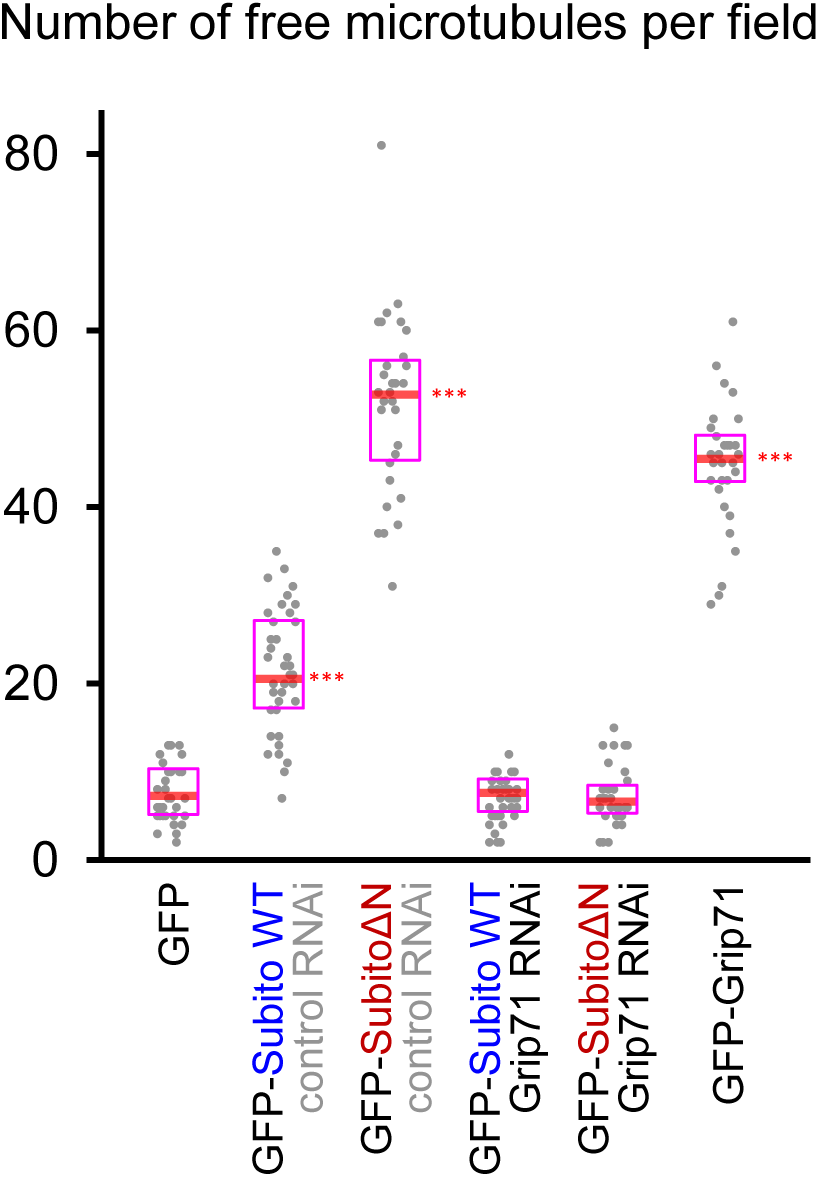
SubitoΔN beads induce microtubule nucleation rather than simply capture microtubules. The number of free microtubules observed per field (51 µm × 102 µm) in the *in vitro* microtubule nucleation assay shown in Figure 5 using immunoprecipitated beads and pure α/β-tubulin dimer (35, 40, 30, 39, 35, 34 fields). The median is indicated by the central line, and the second and third quartiles are indicated by the box. *** indicates a significant difference from wild type (p<0.001). The graph shows the data from one of two repeated experiments, both of which provided similar results. A solution including GFP-SubitoΔN beads had much more free microtubules than one including control GFP beads, confirming that the GFP-SubitoΔN beads induced nucleation of microtubules (some of which were detached from the beads), rather than simply captured microtubules spontaneously nucleated in solution.

